# CHIEF: An Attention-based Ensemble Learning Framework for Functional Protein Design

**DOI:** 10.1101/2025.03.07.641005

**Authors:** Zilong Geng, Yuze Wang, Tingting Liu, Ao Tan, Shuo Wu, Xiaoling Guo, Ruogu li, Xumin Hou, Kun Sun, Lianpin Wu, Qinghua Cui, Lintai Da, Zhiyuan Ma, Honglin Li, Bing Zhang

**Affiliations:** Key Laboratory of Systems Biomedicine, Shanghai Center for Systems Biomedicine, Shanghai Jiao Tong University, Shanghai, China; Engineering Research Center of Techniques and Instruments for Diagnosis and Treatment of Congenital Heart Disease, Institute for Developmental and Regenerative Medicine, Xin Hua Hospital, Shanghai Jiao Tong University, Shanghai, China; Basic Medical Research Center, the Second Affiliated Hospital and Yuying Children’s Hospital of Wenzhou Medical University, Wenzhou, Zhejiang, China; Cardiovascular Department, Shanghai Chest Hospital, Shanghai Jiao Tong University, Shanghai, China; Department of Biomedical Informatics, State Key Laboratory of Vascular Homeostasis and Remodeling, School of Basic Medical Sciences, Peking University, Beijing, China; Department of Electronic Engineering, Tsinghua University, Beijing, China; Shanghai Key Laboratory of New Drug Design School of Pharmacy, Shanghai, China

**Author notes:** Corresponding authors: E-mail addresses (Bing Zhang). These authors contributed equally (Zilong Geng, Yuze Wang, Tingting Liu).

## Abstract

Protein de novo design is a longstanding challenge in biological field due to the multifaceted physicochemical properties and complexities of proteins. Several deep learning models have emerged for protein *de novo* design. However, each model exhibits distinct strengths and limitations because of the differences in training strategies or neural network architectures. Therefore, improving the effectiveness of protein design, particularly for functional protein, requires further improvement. To tackle this challenge, we developed CHIEF (Chimera Ensemble Inverse Folding), an attention-based ensemble framework that integrates five pretrained base models (ProteinMPNN-vanilla, ProteinMPNN-soluble, ESM-IF, Frame2seq and PiFold) to leverage their complementary strengths and mitigate limitations. Compared with the base models, CHIEF significantly improved the sequence recovery rate by 16.6-28.0%, while reducing the prediction perplexity by 22.7-34.6%. CHIEF also exhibited superior protein-designing capacity in variable sequence- and structure-based metrics and in large and complex proteins. CHIEF dynamically captured the semantic information from base models in a context-dependent manner and was minimally affected by the ablation of each base model. More importantly, CHIEF demonstrates real-world applicability to design functional malate dehydrogenase (MDH), achieving a 100% success rate. In summary, our study develops an ensemble deep learning model that improves the efficacy of protein sequence design, and will be a valuable platform for protein engineering and drug development.

## Introduction

The design of functional proteins with desired properties has broad applications in enzyme engineering, drug development and synthetic biology. However, the immense complexity of protein sequence space makes this mission extraordinarily difficult. For example, the theoretical sequence space for a typical small protein with 100 amino acids, is 20^100^, which far exceeds the computing power currently available. The current protein engineering approaches, which rely on the iterative mutagenesis and screening, are inherently limited by their low efficiency and high failure rate^1,2^. For example, random mutagenesis often leads to the loss of activities or functions in over 70% of cases ^3,4^. Thus, there is a compulsive need for more efficient and more innovative strategies in protein design.

The current advancements in deep learning have enabled many complex applications in protein engineering. Deep learning has substantially improved the accuracy of protein structure prediction^5-8^. Conversely, people have also developed several deep neural network models such as ProteinMPNN, ESM-IF, Frame2seq and PiFold that achieve substantial advancements in protein de novo design ^9-17^. These models are built on distinct deep learning architecture and trained with divergent strategies and datasets. ProteinMPNN employs a message-passing neural network to aggregates information from neighboring nodes and achieves a sequence recovery rate of about 49%^9^. ProteinMPNN-vanilla, a derived model of ProteinMPNN excels in designing multimeric proteins due to the enforced training on multi-chain datasets. Another derived model of ProteinMPNN is ProteinMPNN-soluble that is optimized to design the soluble proteins by further training on soluble protein subsets. Unlike ProteinMPNN, ESM-IF employs the GVP-GNN layers to extract geometric features and a generic autoregressive transformer as decoder to capture the semantic information of sequences ^10^. Frame2seq, a structurally conditioned masked language model, uses regularized Invariant Point Attention (IPA) layers to improve the ability to capture protein structural information ^11^. In order to extract more comprehensive features, PiFold introduces a residue perceptron to incorporate learnable virtual atoms^12^. These models exhibit distinct strengths and limitations in different protein-designing practices^18,19^. Therefore, a holistic framework would further improve the performance by leveraging the strengths of these models in different aspects while avert the limitations if it could integrate them well^20,21^.

Ensemble learning, a machine learning technique that combines multiple base learners to improve performance, offers a natural solution for protein design challenges^22^. Ensemble methods have been shown to reduce overfitting and improve model diversity, providing more robust solutions for complex biological problems ^23-25^. However, conventional ensemble methods such as majority voting or weighted voting fail to adapt to the dynamic nature of protein sequences, often assigning fixed weights to each model, which limits their ability to fully capture the complexity of protein sequence design.

In this study, we developed a chimera ensemble inverse folding (CHIEF) model, an attention-based ensemble framework that was able to dynamically and context-specifically integrate the pretrained base models. CHIEF increased the sequence recovery rate to 63.1% in CATH4.2 dataset and 61.8% in PDB dataset and prominently reduced the perplexity. CHIEF also outperformed the base models in variable sequence and structural metrics and in protein variant effect evaluation. As such, CHIEF demonstrates a better ability to design long and multimeric protein complex. We used CHIEF to redesign malate dehydrogenase (MDH) and achieved a success rate of 100%, and the maximal activity of CHIEF-designed MDH was comparable to the wild-type enzymes. In summary, this study provides a new framework to integrate the heterogenous large inverse folding models which exhibits prominently improved performance in protein de novo design.

## Results

### CHIEF improved the protein inverse folding performance

The current pretrained models had distinctive merits and limitations in protein design. To leverage the strengths and mitigate the limitations of these models, we proposed an ensemble learning framework. We chose five well-tested heterogeneous inverse folding models ProteinMPNN-vanilla, ProteinMPNN-soluble, ESM-IF, Frame2seq and PiFold as base models (Fig.1A). To unify the initial contributions from each model, we standardized their outputs into a 21-dimensional probability distribution corresponding to 20 kinds of amino acids and a missing token. Before integrating these probability distributions, we used a sinusoidal positional encoding to tag each amino acid residue with a relative position which would enhance the model’s ability to recognize long-range interactions. The sequence distributions generated from each individual base model were integrated within an ensemble transformer encoder with a self-attention hidden layer and feed-forward neural network where the contribution of each base model was weighted according to the sequence context (Fig.1A). The outputs of the ensemble transformer were the predicted probability distributions over 20 possible tokens for each residue, which were generated through a softmax layer. We trained the ensemble transformer model with the PDB protein dataset, 24,350 proteins (21,359 training, 1,462 testing, 1,529 validation) including 15,991 multi-chain protein structures in order to better capture the structure of multi-unit protein and protein complex. After 200 rounds of training, the cross-entropy loss between the designed and the true sequence distribution was converged to 1.197, and this ensemble learning framework with the pretained components was thereby designated as CHIEF (Chimera Ensemble Inverse Folding).

**Figure 1.**
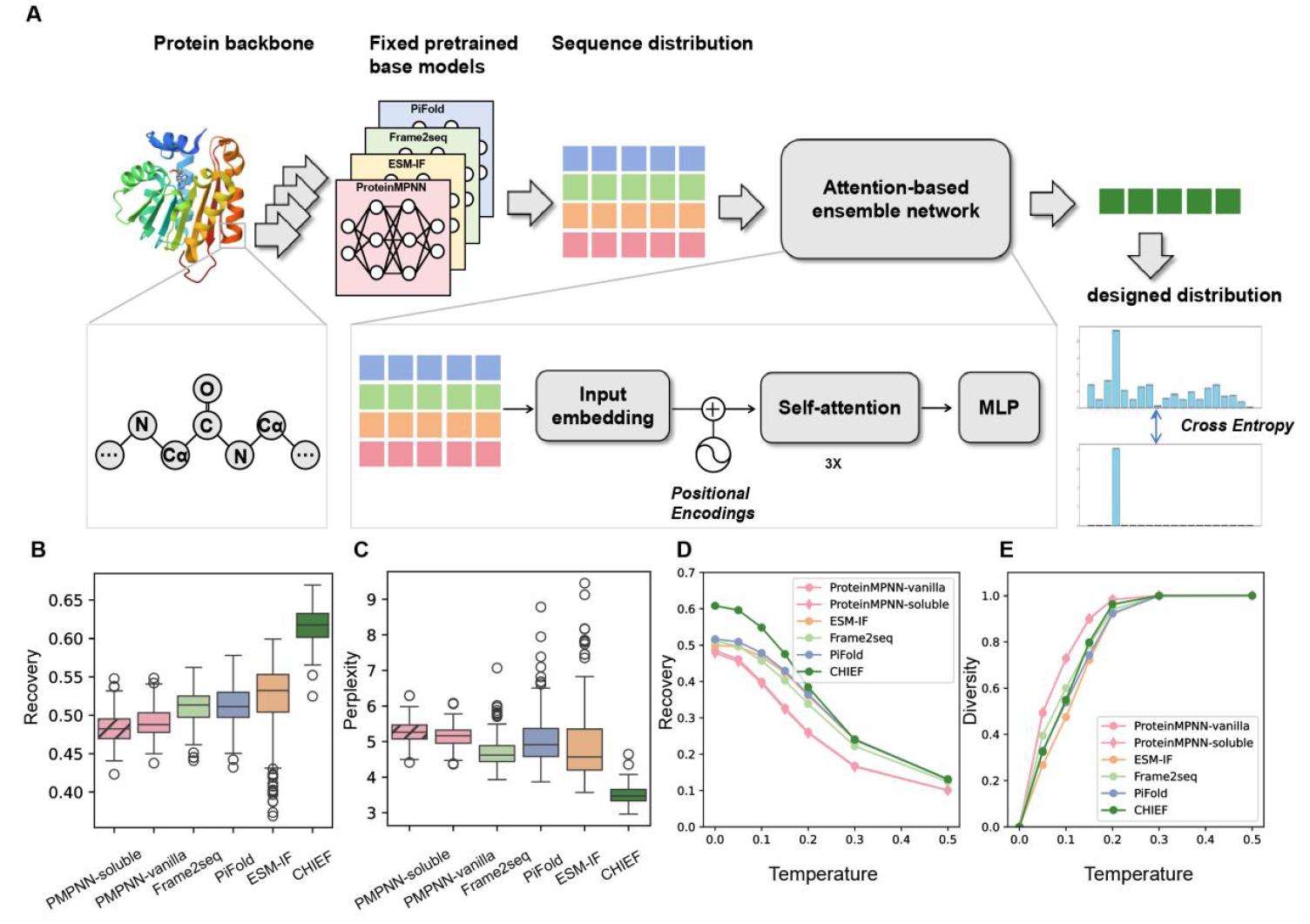
The architecture of CHIEF framework and performance. **A**. The architecture of CHIEF. CHIEF ensemble five inverse folding models: ProteinMPNN-vanilla, ProteinMPNN-soluble, PiFold, Frame2seq, ESM-IF. The inputs are the experimentally validated protein structures from PDB database and augmented with positional encoding before subjected to transformer encoder. Then the hidden representations were fed into MLP, which refines the learned representations and maps them to the final sequence distribution space. **B**. Native sequence recovery rates of base models and CHIEF on testing proteins. **C**. CHIEF has lower prediction perplexity comparing to base models. **D**. Sequence recovery rates in different sampling temperatures. **E**. Sequence diversity of model-generated sequences in different sampling temperatures.

On the test set, CHIEF achieved an average sequence recovery rate of 61.8% which was 16.6% to 28.0% higher than any base model (Fig.1B). As such, CHIEF reduced the perplexity of sequence prediction to 3.4 which was substantially lower than any base model (22.7%-34.6%) (Fig.1C). To further benchmark CHIEF, we examined CHIEF with two other protein datasets TS50 and TS500 commonly used for validation ^26,27^. The CHIEF model achieved average sequence recovery of 70.6% with a perplexity of 2.6 in TS50 (Fig.S1A,S1B), 71.2% and 2.5 respectively in TS500 (Fig.S1C,S1D), which were also better than any base model. Additionally, we tested it on single-chain protein dataset from CATH4.2, where CHIEF achieved a sequence recovery rate of 63.1%. These results demonstrate the generalization of CHIEF’s superior design ability. The model with a high sequence recovery rate usually tends to sacrifice the sequence diversity ^19^, whereas, CHIEF had comparable sequence diversity of designed proteins to base models (Fig.1D,1E). Moreover, to better understand the superiority of CHIEF, we developed two models based on two typical ensemble models, simple averaging and model-based weighted voting, and used them to integrate the same five base models. The sequence recovery rates of averaging and voting models were 56.3% and 59.2%, respectively, which were lower than CHIEF, indicating the better performance of our self-attention learning framework in generating de novo proteins (Fig.S1E,S1F).

### Evaluating the properties of CHIEF-designed proteins

Proteins possess variable sequence and structural properties. To further benchmark the performance of CHIEF, we randomly selected 30 proteins (Tab.S1) with various lengths in different protein categories. For each protein object, 100 sequences designed by each model at a sampling temperature of 0.15 were subjected to sequence and structure assessments (Fig.2A). For sequence-based assessments, we selected two metrics evaluating the homology alignment: BLOSUM62, the substitution matrix score ^28^ and DEDAL^29^, a homology score model; and two metrics given by ESM2 evaluating evolutionary relevance: WTMP, the wild-type marginal probability score and MMP, the masked marginal probability score (Fig.2B). CHIEF demonstrated significant improvements over the base models in homology-based metrics, scoring 38.8% and 28.0% higher on BLOSUM62 substitution matrix and DEDAL model, respectively. CHIEF also had the robust performance on the evolutionary metrics, MMP and WTMP ^30^. Moreover, CHIEF exhibited the lowest sampling variance indicating it’s better consistency in evolutionary prediction (Fig.2B).

**Figure 2.**
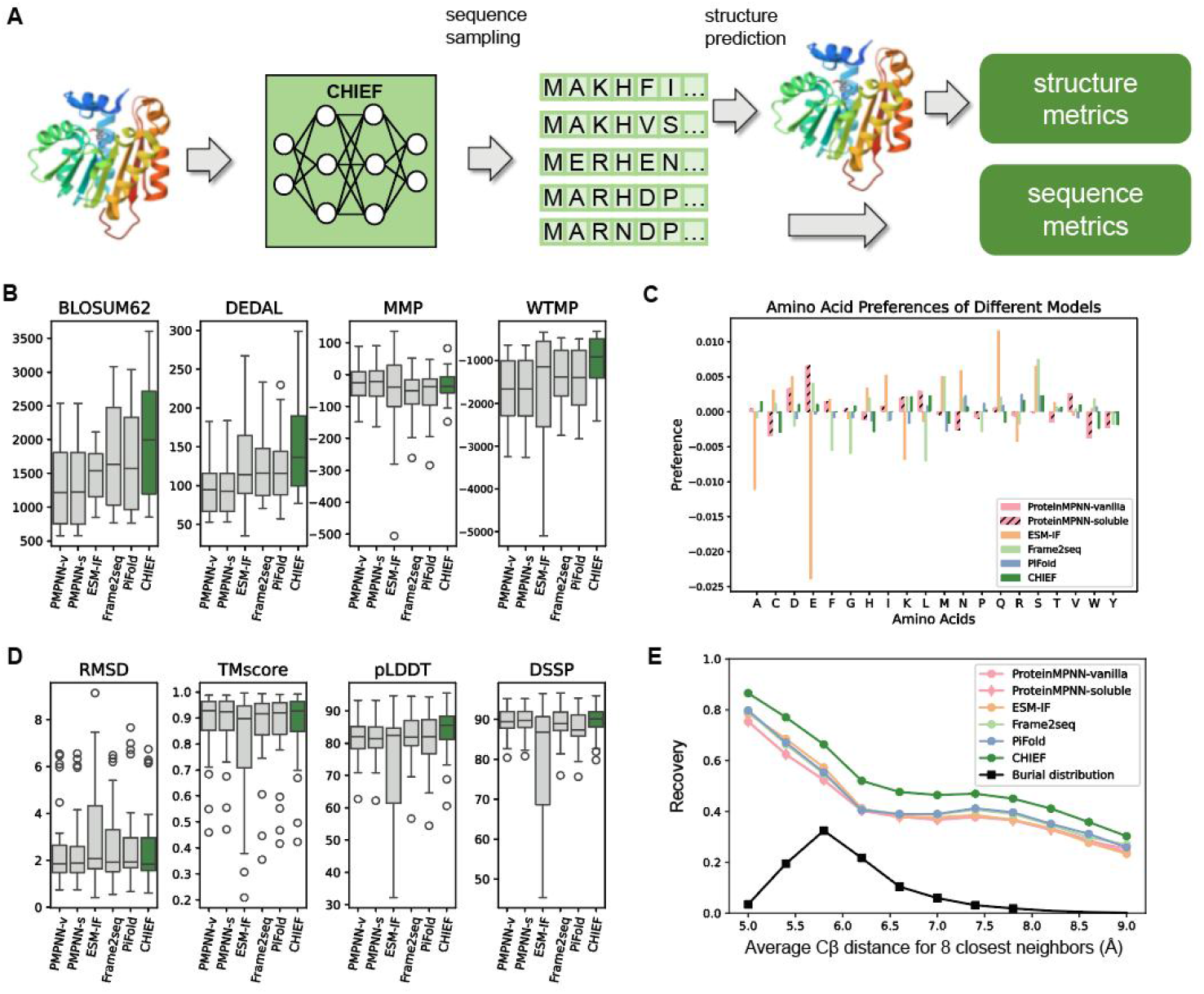
Evaluation of Protein Sequences and Structural Features Designed by CHIEF. **A**.The benchmark workflow. Sequences yielded by the de novo design models and corresponding structures predicted using ESMfold were evaluated by a set of sequence-based and structure-based metrics. **B**. Benchmark results from the sequence metrics including BLOSUM62 substitution matrix scores, DEDAL scores, ESM2 masked marginal probability, and wild-type marginal probability. **C**. Preferences of different models in amino acid usage. **D**. Benchmark results from the structure-based metrics including RMSD and TM-score calculated by comparing the wild-type and predicted structures, pLDDT of the predicted structures, and DSSP evaluating the secondary structure recovery. **E**. Sequence recovery of amino acids with diverse burial rate indicated by average Cβ distance for eight closest neighbors (Å).

The amino acid preferences of inverse folding models may lead to residue prediction biases. We assessed this possible compositional bias by comparing the amino acid frequency of CHIEF-predicted sequences to the frequency of actual sequences in the test set. ProteinMPNN, ESM-IF and Frame2seq demonstrated the apparent amino acid bias. For example, ESM-IF had strong biased usage in N, Q, S, and no usage in A, E and K. ProteinMPNN had biases in D and E, and no usage in C and W, while Frame2seq had biases in M and S and no usages in F, G and L. By contrast, compared to the natural sequences, CHIEF and PiFold exhibited the lowest compositional biases across almost all amino acid types indicating they had more legitimate usages of amino acids (Fig.2C).

Next, we evaluated whether CHIEF-generated sequences could correctly fold. We first employed ESMfold to predict the structures of the designed sequences and used the root mean square deviation (RMSD) and TM-score to quantify the global structural similarity. Among all six tested models, ESM-IF had the highest RMSD and lowest TM score and the largest variance, which was possibly attributed to its suboptimal performance in designing long proteins (Fig.S2). CHIEF had comparable or even better performance in terms of RMSD and TM-score than any base model (Fig.2D). Furthermore, we used pLDDT (predicted local distance difference test) to evaluate the changes in local geometric structure and DSSP to assess the folding accuracy of the secondary structure. Similar to RMSD and TM-Score, ESM-IF scored the lowest in DSSP and exhibited the largest variances in both benchmark indexes. In contrast, CHIEF achieved the best average score in both indexes, and the variances were similar or even lower which indicated the designed proteins by CHIEF had more acurate secondary structure compared to the ones generated by base models.

Another evaluation metric for protein folding is the precision of residue burial within the protein core, a critical factor for protein thermodynamic stability. We calculated the recovery rate of charged residues with varying levels of burial in the proteins from PDB test set and found that compared to other models, CHIEF demonstrated the highest sequence recovery across all different burial levels (Fig.2E), though the distribution of charged residues was similar (Fig.S3). These results together indicated CHIEF had more precise sequence prediction regardless of the residue burial rate.

### Evaluating the functionality of CHIEF-designed proteins

To evaluate the ability of CHIEF to design functional proteins, we assessed the variant effects predicted by CHIEF and base models in deep mutational scanning (DMS) substitution datasets from ProteinGym^18^, which incorporates a large amount of experimentally-tested amino acid substitutions and provides the measurements of fitness changes (Fig.3A). To evaluate the overall precision and efficacy of prediction, we employed Spearman correlation, Area under the curve (AUC), and Matthews correlation coefficient (MCC). The average score of ESM-IF was the highest in all three test indexes while the ProteinMPNN models ranked the lowest, which was consistent with the high performance of ESM-IF in variant effect prediction as previously reported ^18^. CHIEF ranked in the middle and was comparable with the other two base models of Frame2seq and PiFold. We also used normalized discounted cumulative gain and top 10% recall metrics to further benchmark their sensitivity in identifying high-fitness sequences. In these evaluating metrics, CHIEF demonstrated the advantages with a NDCG score of 0.75 and a Top 10% Recall score of 0.22 that were all higher than the base models including ESM-IF (Fig.3B).

**Figure 3.**
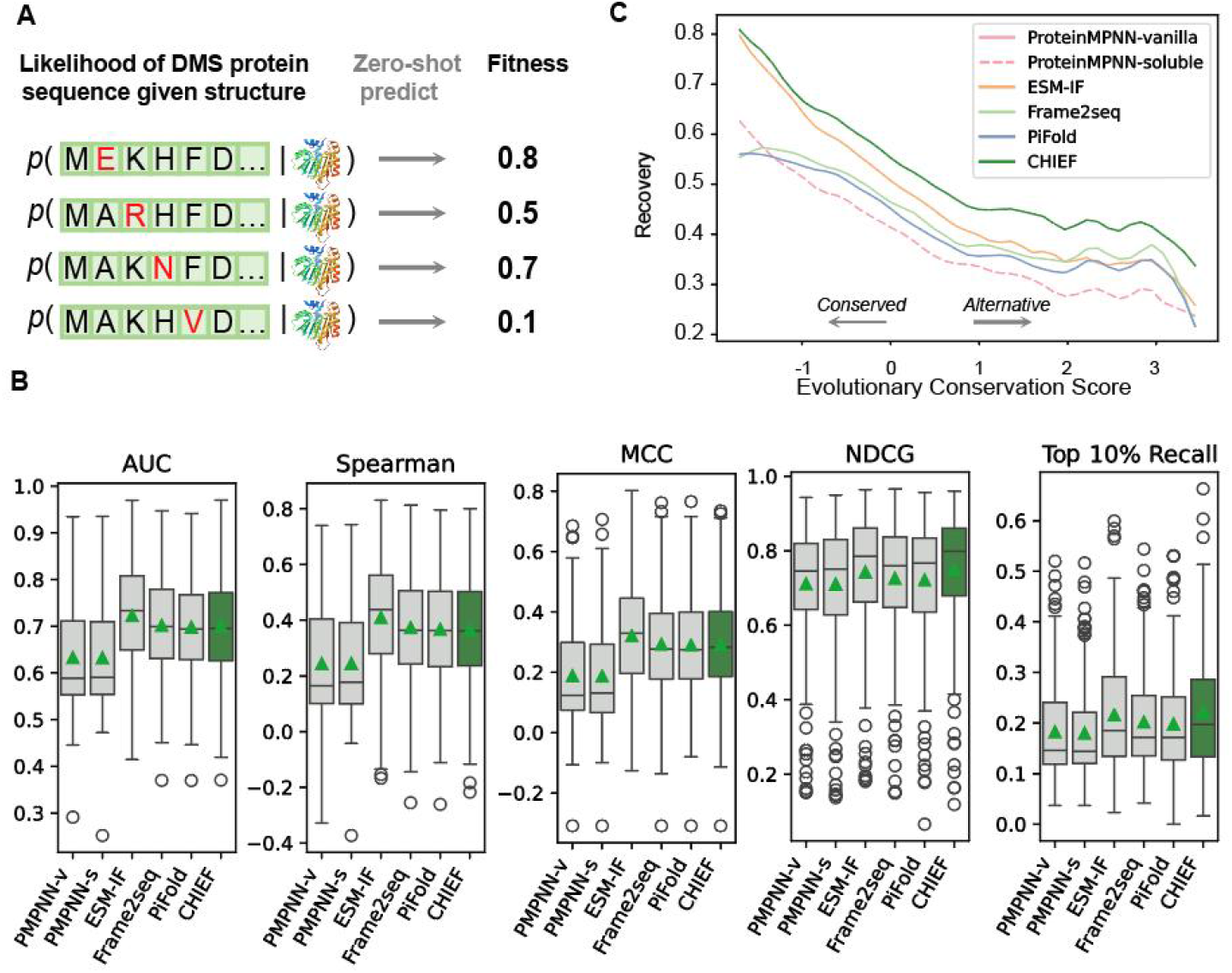
Assessment of the Functional Properties of CHIEF-Designed Proteins. **A**. Evaluating the potent of protein design models that predict variant effects. The benchmarks of pre-verified variant effect were collected from ProteinGym. The likelihood of mutated sequence in the predicted sequences collection representing the mutation effect (fitness) predicted by the protein design models. **B**. Evaluating metrics for variant effect. For all variant effect: AUC: area under curve; Spearman: Spearman’s correlation coefficient; MCC: Matthews correlation coefficient; NDCG: normalized discounted cumulative gain; Top 10% recall: recall the top 10% of variants. **C**. Sequence recovery of amino acid with different conservation score calculated from the CONSURF database.

Evolutionarily conserved amino acid residues have crucial roles in protein folding and functionality. We therefore further assessed the ability of CHIEF and base models in recovering conserved amino acid residues (Fig.3C). For residues with high conservation, CHIEF had a comparable or slightly higher recovery rate than ESM-IF but significantly higher than other models. For residues with low conservation, the recovery rate of ESM-IF and other models declined quickly; however, CHIEF retained its superior ability.

In summary, these results indicate that CHIEF possesses the superior ability to predict variant effects and conservation.

### Evaluating the ability of the CHIEF model in designing long and complex proteins

Natural proteins vary in length and the artificial design for long proteins faces significant challenge due to their much larger sequence space and complex structures. We evaluated the ability of each model in designing proteins with different lengths in the test set. As expected, the sequence recovery declines when the sequence length is greater than 400 amino acids. ESM-IF had high sequence recovery rates for short proteins but declined the most significantly when the length was beyond 400 amino acids, and even lower than 40%, possibly because it was built on an autoregressive model that depends on the sequence orders. ProteinMPNN-soluble and ProteinMPNN-vanilla did not decline very significantly along with increased length, but the recovery rate was the lowest under most conditions. In sharp contrast, CHIEF demonstrated the highest recovery rate, higher than 55% in all tested length windows even longer than 800 amino acids, which indicated the robustness of CHIEF for long protein sequence design (Fig.4A).

**Figure 4.**
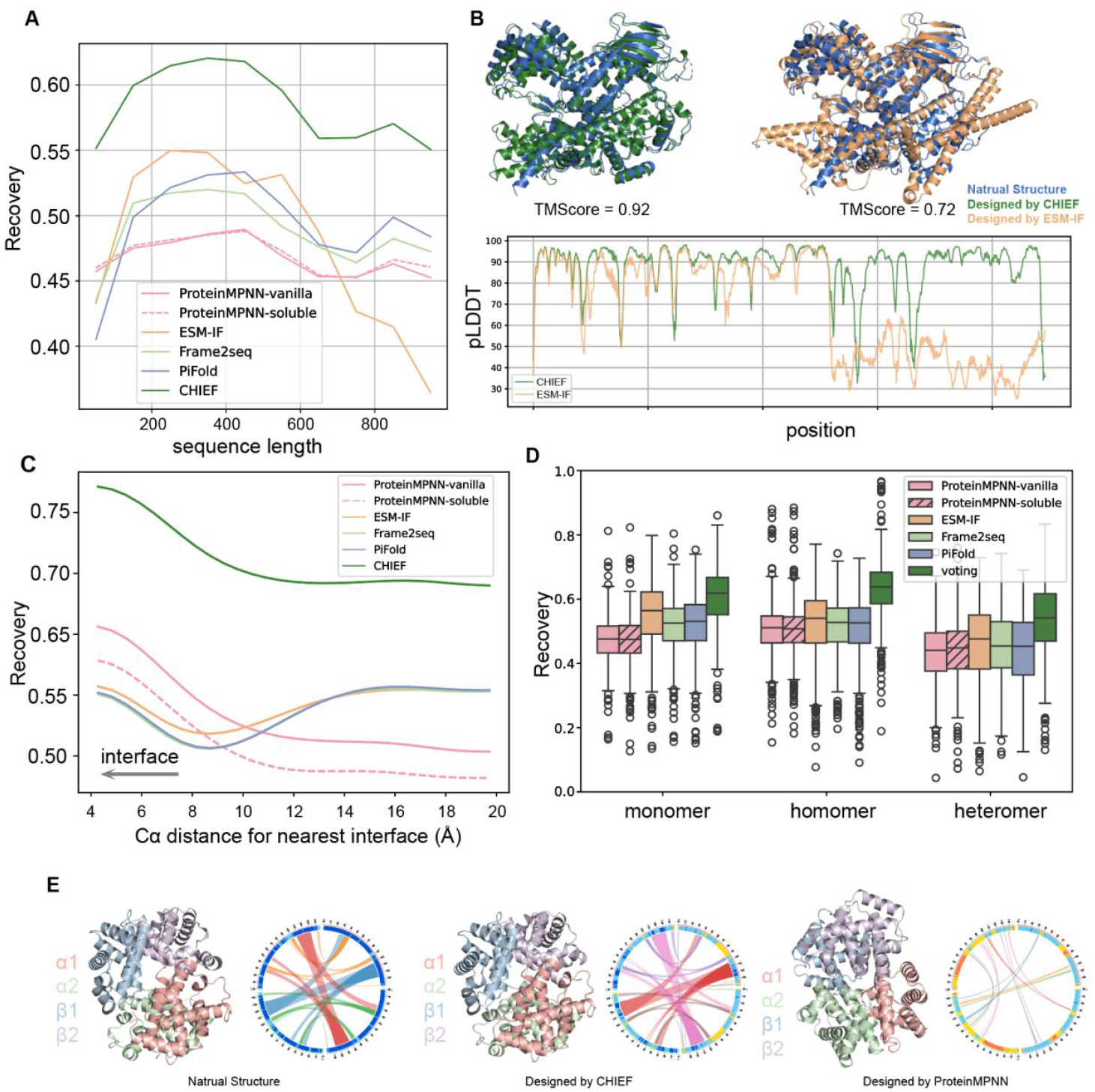
CHIEF facilitates the design of long proteins and protein complexes. **A**. Sequence recovery of the proteins with varying lengths. **B**. The structural examples of Cas13d designed by CHIEF or ESM-IF respectively which were predicted by AlphaFold3. **C**. Sequence recovery rates at the interfaces of protein complexes indicated by Cα distance for nearest interface (Å). **D**. Sequence recovery of monomers, homomers and heteromers from PDB database. **E**. The interface interactions within wild-type, CHIEF-designed and ProteinMPNN-designed hemoglobin tetramer. CHIEF rather ProteinMPNN-designed hemoglobin has similar interactions to wild-type.

The design of new CRISPR RNA nuclease Cas13d served as an example which consisted of 954 amino acids and had been well-studied in our previous research^31^. The results demonstrated that a new Cas13d designed by CHIEF, sampled highly resembled the wild-type Cas13d predicted by AlphaFold3, with a TM-score of 0.92, while the TM-score for the ESM-IF-designed Cas13d was only 0.72 (Fig.4B). The large difference mainly occurred at the C-terminal region of Cas13d beyond the 500th amino acid, where the pLDDT score of the ESM-IF-designed Cas13d was 45.3, but it increased to 86.0 in the CHIEF-designed Cas13d. These results reinforced the notion that CHIEF had a superior competence in designing large proteins.

Proteins often consist of multiple units or function as complexes. Therefore, we further evaluated the ability of CHIEF to design multimeric proteins using a PDB test set with 586 homomers and 423 heteromers^9^. The overall sequence recoveries of CHIEF across variable Ca distances at the protein-protein interface was significantly higher than all base models (Fig.4C). At the closest interface regions (Cα distance=4 Å), the average sequence recovery reached 72.4% (Fig.4D). The second-best model was ProteinMPNN-vanilla, specifically trained on multimers dataset, the sequence recovery rate of it was 60.9%, still significantly lower than CHIEF. We also calculated the sequence recovery rate for protein homomers and heteromers separately. CHIEF achieved 63.2% and 53.2%, respectively, which were all higher than base models as well. To further verify these results, we deployed CHIEF to design human hemoglobin tetramers containing two α and two β subunits and compared them with ProteinMPNN-vanilla. The average TM-score of CHIEF-designed hemoglobin is 0.92 and the average sequence recovery rate of interface residues is 68.8%. Both were higher than the hemoglobins designed by ProteinMPNN that had a TM score of 0.88 and recovery rate 56.9%, respectively. We further utilized the Alpha-bridge to visualize the interactions between the subunits and found 8 out of 10 CHIEF-designed sequences had similar interaction patterns to nature hemoglobin (Fig.S4), and in sharp contrast, no sequence designed by ProteinMPNN-vanilla recapitulated the interactions (Fig.4E). These results further validated the ability of CHIEF in designing multimeric protein complexes.

### CHIEF dynamically captured the implicit semantic information from base models

To explore the contribution of each individual base model to CHIEF, we conducted an ablation study: we removed the base model one by one from the CHIEF ensemble and then evaluated the results (Fig.5A). Among the five base models, ESM-IF demonstrated the highest impact on sequence recovery and prediction perplexity, but overall, the exclusion of a single base model had minor effect on CHIEF’s performance which indicated that CHIEF effectively integrated all base models rather than relying on individual models (Fig.5B). To further benchmark CHIEF’s ensemble ability in the sequence and structural metrics we tested above, we redesigned the same set of 30 proteins as described above with the ablated CHIEF models by following the same procedure. In consistence, the exclusion of any single base model did not substantially affect CHIEF’s ability in protein design though in some sequence metrics ESM-IF had some relatively high impacts (Fig.S5). These results further verified the fact that CHIEF was able to efficiently integrate all base models.

**Figure 5.**
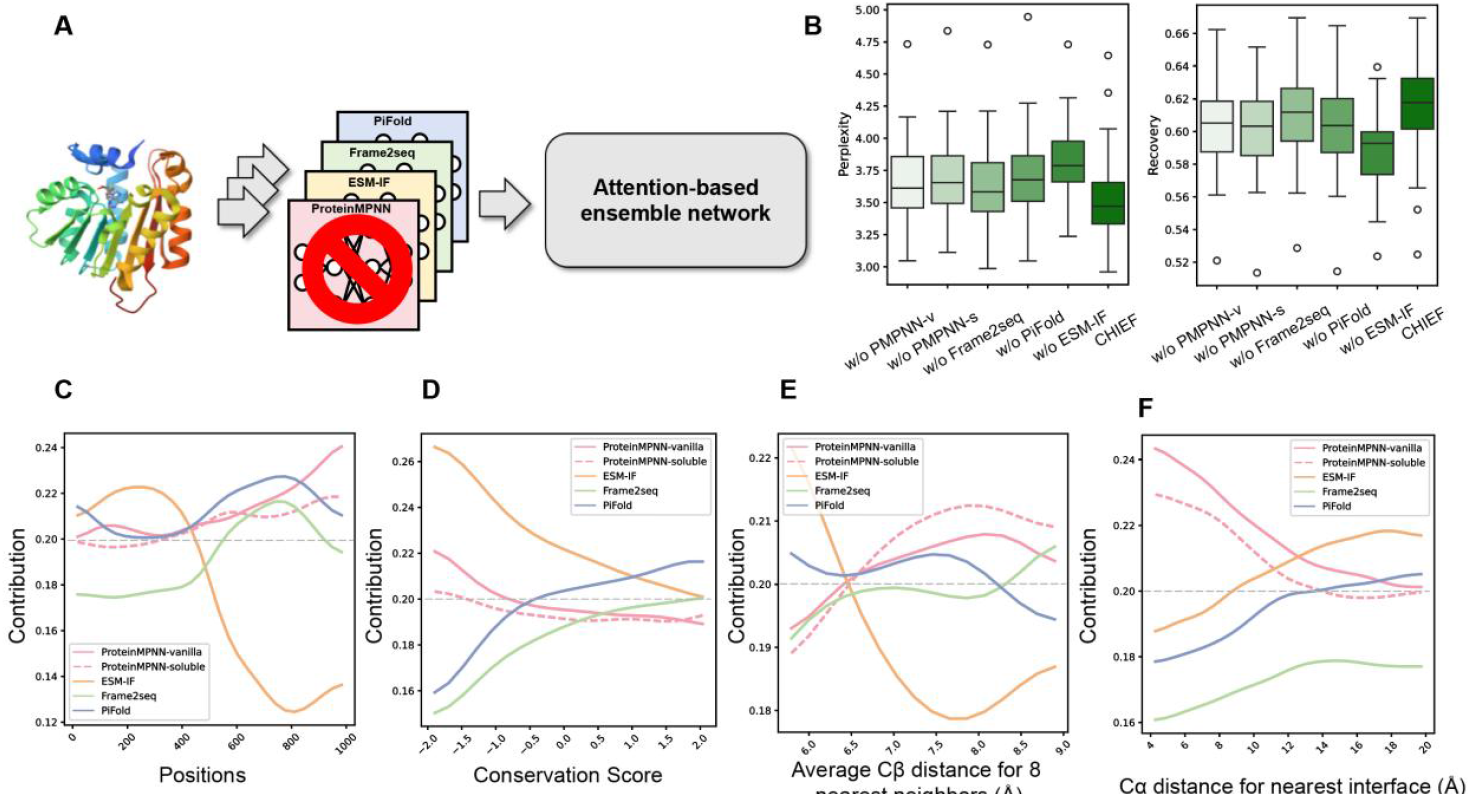
An ablation study evaluating the contribution of the base models. **A**. The scheme of the ablation experiments. Each of five base model was ablated once a time. **B**. The sequence recovery and perplexity of CHIEF after ablation of the individual base model. **C**. The contribution of each base model at different amino acid residue. The gray dashed line represents the baseline (0.2) of the average voting strategy. **D**. The contribution of each base model at residues with varying degrees of evolutionary conservation. **E**. The contribution of each base model at residues with different levels of solvent accessibility. **F**. The contribution of each base model at protein-protein interaction interfaces.

CHIEF is an attention-based sequence ensemble model which implied that CHIEF could implicitly capture the semantic information in addition to structure. We used a least squares regression approach to evaluate the semantic contribution of each base model to CHIEF. We first assessed the contributions of base models across residue positions. For the first 500 residues, ESM-IF demonstrated higher impact, but the influence quickly declined (Fig.5C), which was consistent with the fact that ESM-IF was trained with the proteins only shorter than 500 amino acids. For the residues surpassing 500, the major contributors were shifted to the other four models especially ProteinMPNN-vanilla that utilized a random-order decoding to avoid the limitation imposed by sequence length. Next, we evaluated the contributions of the base models on conservation (Fig.5D). We found ESM-IF had better performance to identify the highly conserved residues. Consistent with these findings, ESM-IF exhibited the highest contribution to highly conserved residues, whereas Frame2seq and PiFold showed the least.

**Figure 6.**
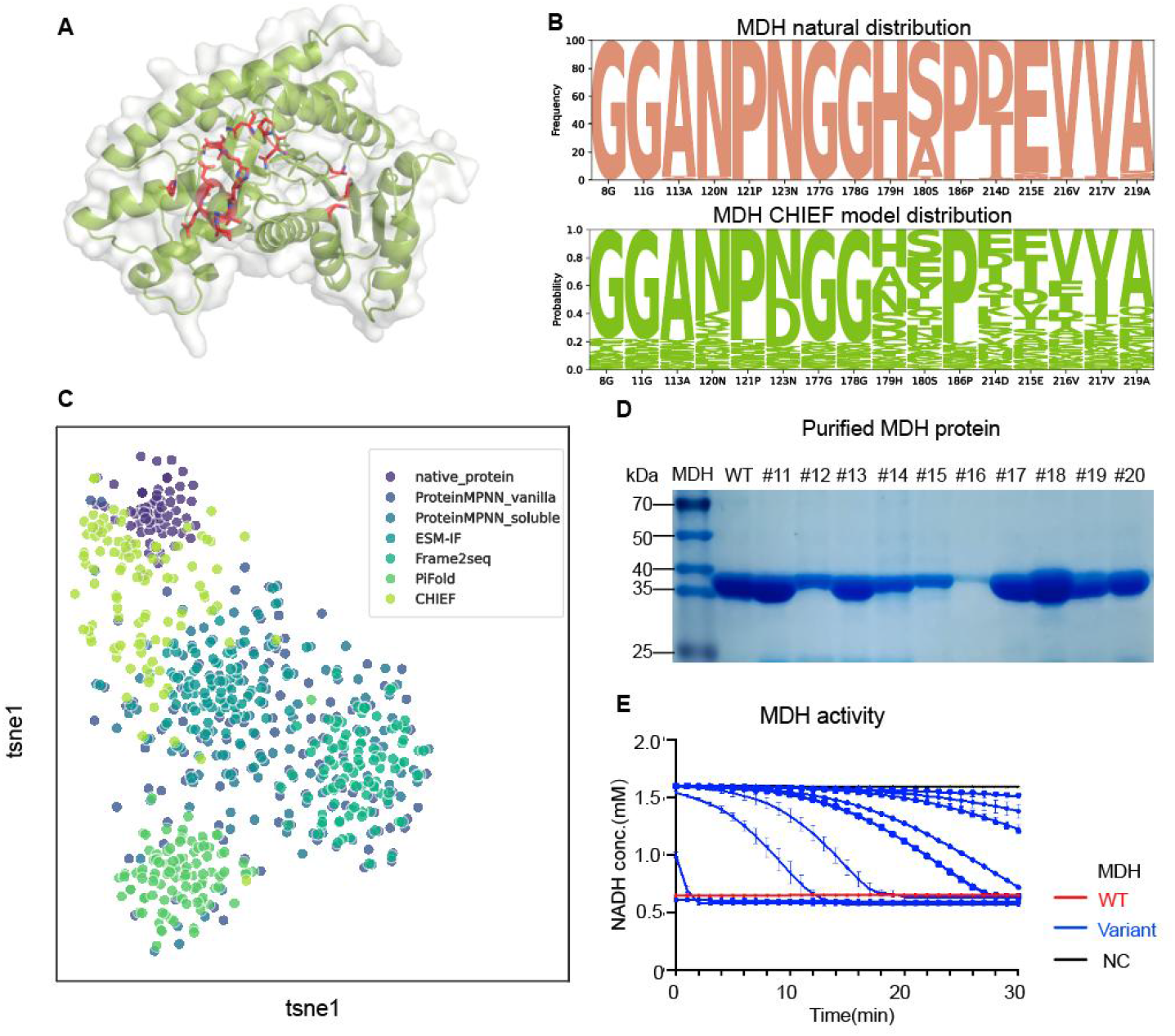
Redesign of Malate Dehydrogenase (MDH) by CHIEF. **A**. MDH three-dimensional structure predicted by AlphaFold2, with key functional residues highlighted in red. **B**. Sequence logo comparison of the natural amino acid distribution and CHIEF-designed amino acid distribution at evolutionarily conserved sites in MDH, illustrating sequence differences and conservation patterns. **C**. t-SNE visualization of the sequence sampling space, showing the distribution of CHIEF-designed MDH sequences in comparison to those generated by other models and native MDH sequences. **D**. SDS-PAGE analysis of purified CHIEF-designed MDH variants, demonstrating successful protein expression. **E**. Enzymatic activity assays comparing CHIEF-designed MDH variants to wild-type MDH (WT), a variant group, and a negative control (NC). NADH consumption over time is plotted, highlighting the functional performance of CHIEF-designed MDH.

The solvent exposure was the third considered assessment. ESM-IF had higher weights for the residues closer to protein core, while ProteinMPNN-soluble contributed more to the amino acid residues nearby the protein surface (Fig.5E). ProteinMPNN-soluble was specifically trained on the soluble proteins and the expertise at capturing the hydrophilic sequences on the protein surface. Finally, we evaluated the contributions of base models on protein-protein interaction and found ProteinMPNN-vanilla and -soluble contributed the most to the residue recovery rate near the protein-protein interface (Fig.5F). It was also in agreement with the fact that ProteinMPNN was trained on multichain proteins^9^. Collectively, these results indicate that CHIEF as ensemble model is able to precisely distilled implicit semantic information embedded in the sequences from the base models and generated a new sequence probability distribution that was optimal for the protein properties and functions.

### Redesign functional malate dehydrogenase with CHIEF

CHIEF demonstrated an improved ability in various metrics to design de novo protein. We next sought to validate it in real experiments. Malate Dehydrogenase (MDH) is a key enzyme in the Krebs cycle catalyzing malate to oxaloacetate^32^. New MDH enzymes had been designed by protein-design models, however, the functionality rate required further improvements^20,33,34^. An AlphaFold-predicted structure (Uniprot: A0A319AA41) was used as a backbone template in this assay (Fig.6A). At a sampling temperature of 0.15, CHIEF generated 1000 sequences with a sequence diversity of 0.28 and naive sequence recovery 64.4% to 70.9%. Their evolutionary conservation pattern closely resembled natural sequences (Fig6.B). Within a sampling space,

CHIEF-generated sequences were apparently closer to the wild-type ones when compared with the ones from base models which suggested a higher similarity between CHIEF-generated sequences and wild-type ones (Fig.6C). We synthesized the 10 sequences generated by CHIEF and validated them by the activity assay. All 10 sequences were expressed in BL21 *E*.*coli* cells, and were able to be purified and resolved in the assay buffer. The activity assay illustrated all designed sequences exhibited observable enzymatic activity, and one sequence show an even comparable activity to the wild-type MDH (Fig.6D&E). These results illustrated a superior ability of CHIEF to generate functional enzymes.

## Discussion

Protein design is a fundamental challenge in biological field due to the infinite sequence space, multi-dimensional traits and diverse functions of proteins^35-38^. The rapid advancement of artificial intelligence makes protein design practical. Recently emerging inverse folding models can generate de novo polypeptides with structures recapitulating their references. However, due to the distinct neural network architectures and training strategies, their performances in diverse protein metrics vary and the ability to design functional protein requires further improvement. Here, we developed a self-attention-based ensemble model CHIEF to integrate multiple available protein-design tools and demonstrated superior performances in versatile scenarios. CHIEF outperformed other models in sequence recovery rate and perplexity. CHIEF dynamically and selectively leveraged the strength of its base models and demonstrated the superior potent in diverse sequence and structure metrics. More importantly, CHIEF was able to design functional MDH and all designed MDH variants exhibited activity. Collectively, through an ensemble learning of diverse pretrained base models, CHIEF provides an improved platform for protein design.

Ensemble learning is a machine-learning paradigm using the idea of “the wisdom of the crowd outperforms the individual” and by integrating multiple models, is able to improve the prediction performance. Ensemble learning has been widely used in data mining, image classification and nature language processing^39,40^. The widely used combination approaches are averaging and weighted voting whose weights assigned for base models are fixed. Therefore, they can’t dynamically and precisely capture the specific merits of each base models and can’t have the performance beyond the linear combinations of base models. To overcome these limitations, we developed an attention-based ensemble model to capture and process the semantic information of output sequences yielded by base models and generate new sequences with optimal functions. The transformer embedded in CHIEF was able to capture the contextual semantics to balance the strengths and limitations of each model under different conditions. In consequences, CHIEF achieved the best outcomes in most sequence and structural scenarios such as sequence recovery rate, prediction perplexity, amino acid burial and sequence and structure similarity etc. The ablation studies further verified it by illustrating the robustness of CHIEF upon the absence of base models. It worth to mention that although ensembled five base models with protein structures as input, CHIEF is able to capture the semantic implication because the transformer in CHIEF perceived and processed sequence distributions, which facilitate the capture of protein functions.

In this study, we only included five inverse folding deep leaning models, but there were other emerging models bearing diverse merits, especially protein language models such as Progen2 and ESM2^41,42^, that better capture the semantic information of proteins and provides more diversity and flexibility for protein design. Extending the current set of base models in CHIEF framework by including language protein models could possibly further increase the model performance. The recent diffusion models are applied to generate the naturally nonexistent proteins and require a conversion of structure to sequence^43^. Considering an excellent performance of CHIEF in structure-to-sequence conversion, linking CHIEF with the diffusion models would increase the accuracy and functionality of designed proteins.

A key feature of CHIEF lies in its ensemble learning strategy. While effective, this approach could be further enhanced by deploying more advanced architectures such as Mixture of Experts (MoE) framework which are prevailing in large-scale AI models^44,45^. In a MoE framework, a greater number of distinctive expert models could be incorporated to better capture and process diverse aspects of protein-related information, including sequence, structure and function. We could directly include the sequence design model such as Progen2 and ESM2 aforementioned in addition to the inverse folding models used in this study to make the framework more extendable. The router mechanism used by MoE will also make the integration of expert models more efficiently and reduce the amount of computation.

In summary, we developed CHIEF as an optimized ensemble framework for inverse folding models which provides a new platform for protein design and engineering.

## Methods

### CHIEF’s architecture and training

The CHIEF model was trained with proteins with experimentally validated three-dimensional structures deposited in Protein Data Bank (PDB).In line with the procedure employed by ProteinMPNN, the dataset was filtered with a maximal 30% sequence identity ensuring the diversity while reducing redundancy. The data were separated into training (23,358 clusters), validation (1,464 clusters), and testing (1,539 clusters) sets to pretrain or evaluate the model performance.

Atoms N, Cα, C, and O in a fixed structural scaffold was served as input. For each base model in the ensemble, the log-probability distributions of amino acid residues were calculated from the input backbone structure. Therefore, output from each base model was a tensor of shape [N, M, 21], where N is the number of residues in the sequence, M is the number of base models, and 21 is 20 standard amino acid types plus a missing token X. These tensors generated by each base model were the concatenated and flattened into a [N, M × 21] tensor in order to integrate the output across multiple models. To capture the positions of amino acids within the sequence, the flattend representations were augmented with sinusoidal positional encodings that provides the spatial information and enable the model to consider the sequence order in the folding and design process.

These outputs were then fed into the CHIEF model’s attention-based ensemble network, which used a transformer architecture^46^. The model employed a 3-layered transformer encoder with a hidden dimension of d_model=128, designed to weigh and combine the outputs from the five base models (ProteinMPNN-vanilla, ProteinMPNN-soluble, ESM-IF, Frame2seq, and PiFold). The attention network contains a Transformer Encoder with three layers and four attention heads. The training objectives were the protein inverse folding formulated as the minimized negative log-likelihood (NLL) loss. The training loss was computed as:

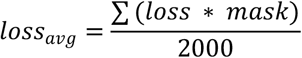

where the mask was the missing or padding positions of amino acid. The Adam optimizer was employed with a learning rate of 0.001, and the training was performed for over 200 epochs using a NVIDIA A100 GPU. All models were trained to converge, and validation loss was monitored during the entire procedure. All the codes were deposited to github.

### The benchmark in sequence and structure metrics

The PDB test set was divided into three categories based on protein length: short (200-500 aa), medium (500-800aa), and long (800-1000 aa) and ten distinct protein chains were randomly selected from each category. For each model, 100 polypeptide sequences were generated at a sampling temperature of 0.15. DEDAL ^29^ scored the homology between the designed and the original sequences. BLOSUM62 calculated the substitution score. esm2_t33_650M_UR50D calculated the masked marginal probability scores and the wild-type marginal probability scores. pLDDT scored the local distance difference generated by ESMFold^30^. TMsalign compute the TM-score and RMSD. DSSP calculate the secondary structure recovery rates^47^. The structural for designed sequence were predicted by ESMFold.

### Benchmarking Inverse Folding Models on the ProteinGym Dataset

To benchmark the mutation effect prediction performance of inverse folding models, we use the substitution dataset from ProteinGym, which provides 2.4M experimentally measured deep mutational scanning (DMS) substitutions data for a wide range of proteins. The dataset consists of protein sequences where single or multiple amino acid substitutions have been introduced, along with corresponding fitness measurements reflecting functional impacts. Since inverse folding models require structural input, missing protein structures were generated using AlphaFold2 (AF2) to ensure that each sequence has an associated structural representation.

For each sequence in the ProteinGym dataset, we compute a likelihood-based score using negative log-likelihood (NLL). Given a target sequence *S*, the inverse folding model predicts a probability distribution over amino acids at each position given structure. The NLL score is computed as:

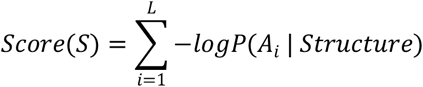

where *P* (*A*_*i*_ | *structure*)is the model-predicted probability of the amino acid at position *i* given structure, and *L* is the length of protein.

### Ablation assay and Interpretability study

To investigate the contribution of each component in CHIEF, we conducted an ablation study by systematically removing individual base models and assessing the impact on performance. For each ablation experiment, we removed one of the five base models from CHIEF while keeping the remaining models unchanged. We then retrained the CHIEF framework using the same training and validation data to ensure fair comparison.

To quantify the contribution of each model at individual amino acid residue positions, we employed least squares fitting and applied the softmax function to normalize the model weights, constraining them within the range of 0 to 1. Subsequently, we used ConSurfDB to extract the conservation scores of amino acid residues for test set^48^.

### Enzyme Expression and Purification of MDH

Plasmid construction was performed using E. coli DH5α, while E. coli BL21 (DE3) was utilized for gene expression. The gene was synthesized into the pET28 expression plasmid. All BL21 (DE3) strains harboring the expression plasmids were cultivated in Luria–Bertani (LB) liquid medium containing 5 g/L yeast extract, 10 g/L tryptone, and 10 g/L NaCl. The cultures were incubated at 37°C with constant shaking at 220 rpm. The fermentation of the engineered strains was carried out in Terrific Broth (TB) medium, consisting of 24 g/L yeast extract, 12 g/L peptone, 4 mL glycerol, 2.31 g/L KH2PO4, and 12.54 g/L K2HPO4, and supplemented with 50 µg/mL kanamycin, in accordance with the screening markers of the expression plasmids.

For shake flask fermentation, the engineered strains containing different expression plasmids were first cultured overnight in 5 mL of LB liquid medium at 37°C. Subsequently, 50 µL of the seed culture was used to inoculate 5 mL of TB medium. The cultures were grown at 37°C until an OD600 of 2-4 was reached. Then, Isopropyl β-D-1-thiogalactopyranoside (IPTG) was added to a final concentration of 0.3 mM, and fermentation was allowed to proceed for an additional 10 hours at 28°C.

### MDH Enzyme Activity Assay

To assess L-MDH activity, 1 mL of fermentation broth was collected and centrifuged at 15,000 × g for 3 minutes at 4°C to collect the cell pellet. The pellet was resuspended in 1 mL of phosphate-buffered saline (PBS). The cell suspensions were lysed via sonication on ice, using 30% power with 5-second pulses and 5-second intervals for a total of 10 minutes, yielding the crude MDH enzyme solution. The total reaction volume was 200 µL, comprising 50 mM PBS (176 µL), 200 mM oxaloacetic acid (2 µL), 200 mM NADH (2 µL), and 20 µL of the diluted crude enzyme solution. The absorbance change at 340 nm was monitored, with readings taken every 10 seconds for 3 minutes. Enzyme activity was calculated using the formula: U/min = [ΔA/min] × [1/ϵ] × 1000, where ϵ = 6.22 L/(mmol·cm).

## Acknowledgements

This work was supported by the National Key Research and Development Program of China (2020YFA0803800, 2020YFA0803802, 2023YFA1800700, and 2023YFA1800702); National Foundation of Distinguished Young Scholar of China (82225006); National Natural Science Foundation of China (NSFC 32200926); the Innovation Program of the Shanghai Municipal Education Commission (2021-01-07-00-02-E00088); SJTU STAR Award (YG2022ZD023 &YG2023QNB13); Science and Technology Innovation Action Plan of Shanghai Science and Technology Commission (24J12800400).

## Author contributions

B.Z. supervised this study. Z.L.G. constructed and trained the model. Z.L.G., and Y.Z.W. were responsible for protein design and bioinformatics analysis. T.T.L., and A.T. constructed the expression vectors and performed experimental validation. B.Z., Z.L.G., Y.Z.W., Z.Y.M., and L.T.D. wrote the manuscript. Z.Y.M., Q.H.C., L.T.D., X.M.H., S.W., X.L.G., R.G.L., and L.P.W. provided insights into study design. Z.Y.M., Q.H.C., and L.T.D. provided guidance on bioinformatics methodology. B.Z., H.L.L., and K.S. reviewed and revised the final manuscript.

## Data and code availability

The training data used in this study can be accessed at https://github.com/dauparas/ProteinMPNN. The code supporting this study is currently being organized and refined for public release. It will be made available online shortly after the finalization of documentation and repository structuring. A link to the repository will be provided upon release.

## Figures and figure legends

**Supplementary Figure 1.**
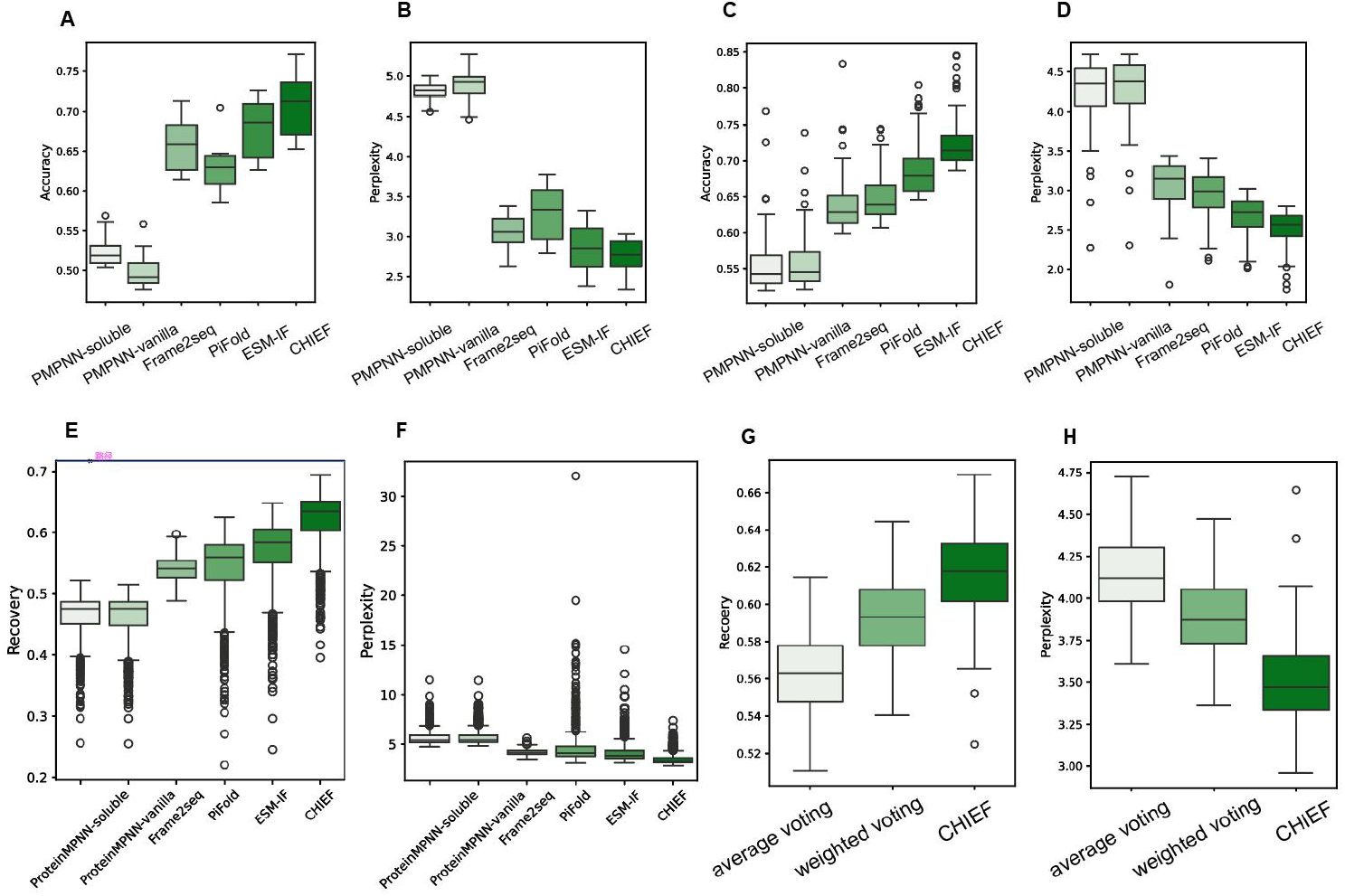
Performance of CHIEF across different datasets and comparison with alternative ensemble methods. **A**. Sequence recovery rate (Q1) of individual base models and CHIEF on the TS50 dataset. **B**. Prediction perplexity (Q1) of individual base models and CHIEF on the TS50 dataset. **C**. Sequence recovery rate (Q1) of individual base models and CHIEF on the TS500 dataset. **D**. Prediction perplexity (Q1) of individual base models and CHIEF on the TS500 dataset. **E**. Sequence recovery rate of individual base models and CHIEF on the CATH 4.2 dataset. **F**. Prediction perplexity of individual base models and CHIEF on the CATH 4.2dataset. **G**. Comparison of sequence recovery rates between CHIEF and two alternative ensemble strategies: mean voting and model-based weighted voting on the PDB test set. **H**. Comparison of prediction perplexity between CHIEF and two alternative ensemble strategies: mean voting and model-based weighted voting on the PDB test set.

**Supplementary Figure 2.**
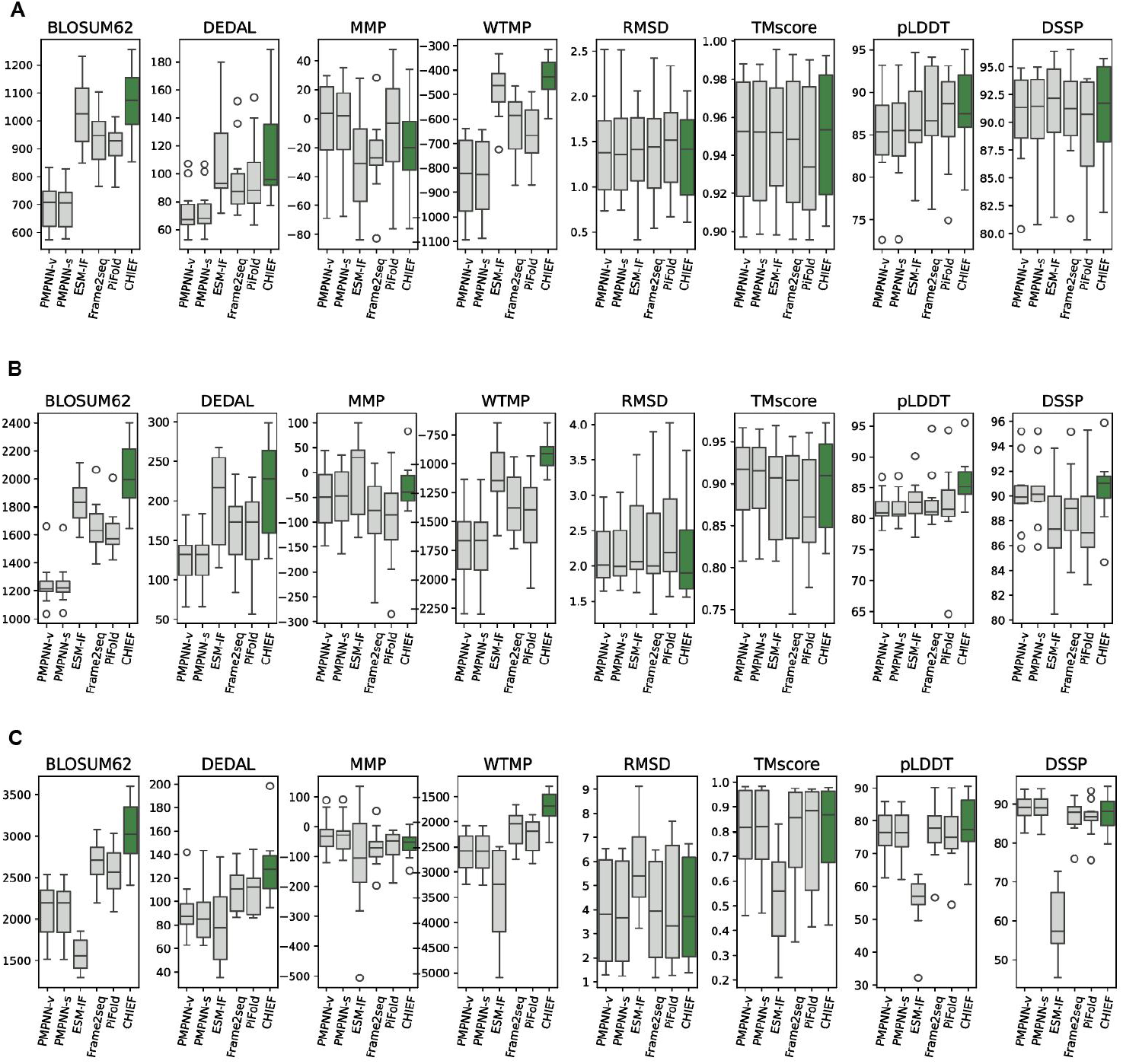
Sequence- and structure-based evaluation metrics of CHIEF in designing proteins of different lengths. **A**. Sequence- and structure-based evaluation metrics of CHIEF in designing proteins with lengths of approximately 300–400 amino acids. Sequence-based metrics include BLOSUM62 substitution matrix scores, DEDAL homology scores, and MMP and WTMP scores derived from ESM2. Designed sequences were structurally predicted using ESMFold, and structure-based metrics include TM-score, RMSD, pLDDT, and secondary structure recovery based on DSSP. **B**. Sequence- and structure-based evaluation metrics of CHIEF in designing proteins with lengths of approximately 500–600 amino acids. **C**.Sequence- and structure-based evaluation metrics of CHIEF in designing proteins with lengths of approximately 900–1000 amino acids.

**Supplementary Figure 3.**
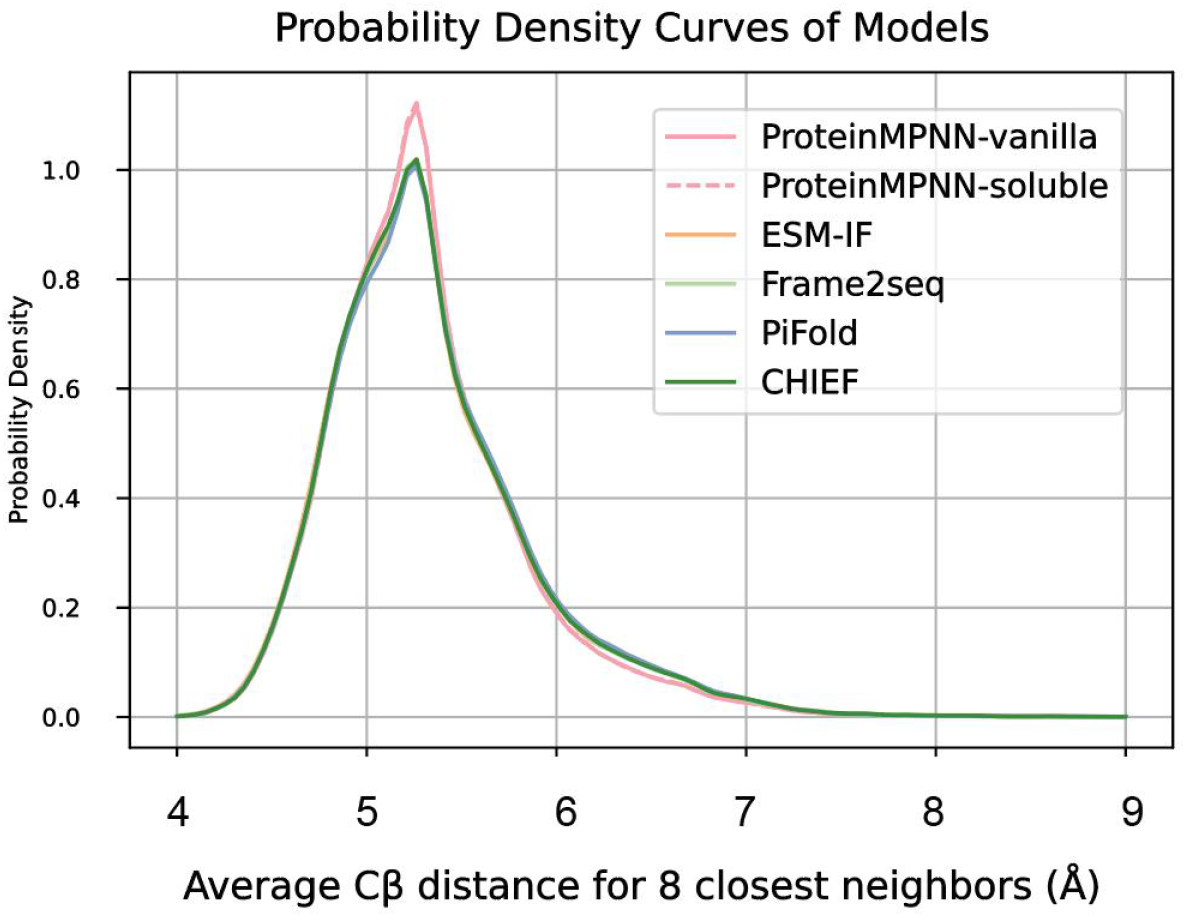
Usage frequency of charged amino acids by different models across varying levels of residue burial.

**Supplementary Figure 4.**
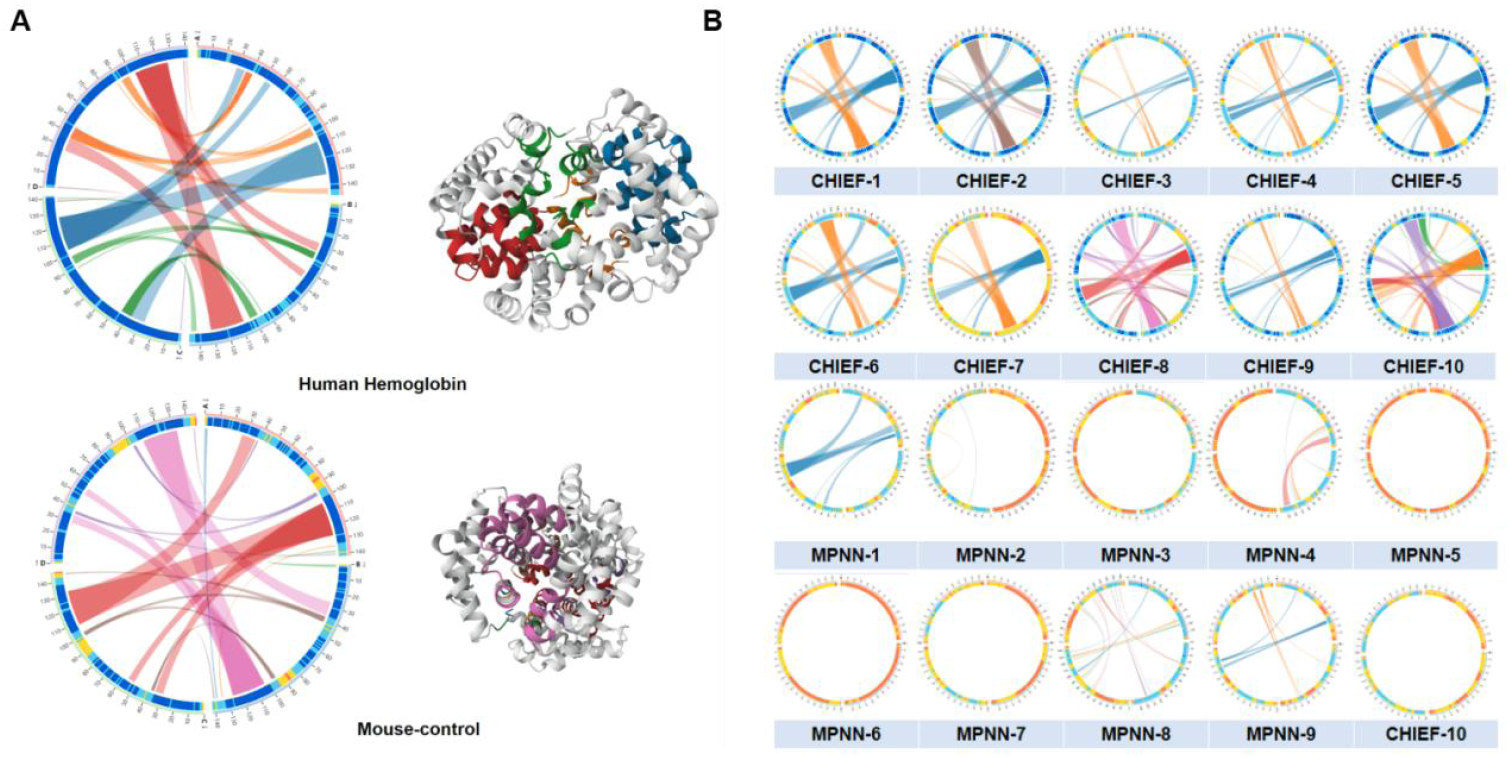
Prediction of subunit interactions in hemoglobin tetramers designed by CHIEF. **A**. Complex structures of human and mouse hemoglobin tetramers predicted by AlphaFold3, along with subunit interaction predictions using Alpha-Bridge. **B**. Predicted subunit interactions for ten sequences designed by CHIEF and ProteinMPNN at a sampling temperature of 0.15. Sequences generated by CHIEF exhibit subunit interaction patterns similar to those observed in native hemoglobin tetramers.

**Supplementary Figure 5.**
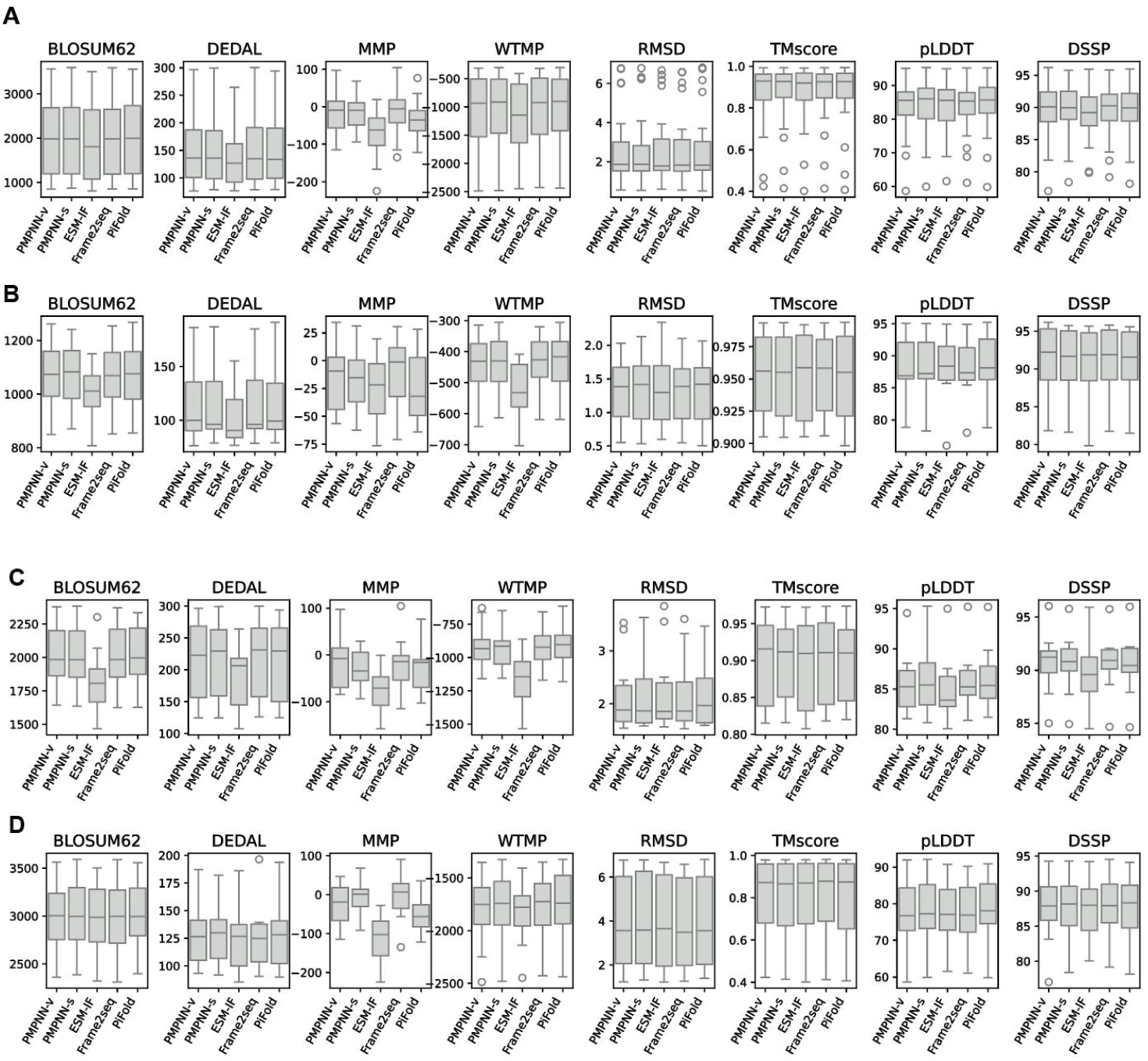
Performance of CHIEF across different metrics after the removal of individual base models. **A**. Average performance of CHIEF across eight metrics after the removal of individual base models in proteins of three different length ranges. **B**. Performance of CHIEF across eight metrics after the removal of individual base models in short proteins (300–400 amino acids). **C**.Performance of CHIEF across eight metrics after the removal of individual base models in medium-length proteins (500–600 amino acids). **D**.Performance of CHIEF across eight metrics after the removal of individual base models in long proteins (900–1000 amino acids).

## Reference

1 Jackel, C., Kast, P. & Hilvert, D. Protein design by directed evolution. Annu Rev Biophys 37, 153–173, doi:10.1146/annurev.biophys.37.032807.125832 (2008).

2 Lutz, S. & Iamurri, S. M. Protein Engineering: Past, Present, and Future. Methods Mol Biol 1685, 1–12, doi:10.1007/978-1-4939-7366-8_1 (2018).

3 Rockah-Shmuel, L., Toth-Petroczy, A. & Tawfik, D. S. Systematic Mapping of Protein Mutational Space by Prolonged Drift Reveals the Deleterious Effects of Seemingly Neutral Mutations. PLoS Comput Biol 11, e1004421, doi:10.1371/journal.pcbi.1004421 (2015).

4 Sarkisyan, K. S. et al. Local fitness landscape of the green fluorescent protein. Nature 533, 397–401, doi:10.1038/nature17995 (2016).

5 Baek, M. et al. Accurate prediction of protein structures and interactions using a three-track neural network. Science 373, 871–876, doi:10.1126/science.abj8754 (2021).

6 Abramson, J. et al. Accurate structure prediction of biomolecular interactions with AlphaFold 3. Nature 630, 493–500, doi:10.1038/s41586-024-07487-w (2024).

7 Jumper, J. et al. Highly accurate protein structure prediction with AlphaFold. Nature 596, 583–589, doi:10.1038/s41586-021-03819-2 (2021).

8 Senior, A. W. et al. Improved protein structure prediction using potentials from deep learning. Nature 577, 706–710, doi:10.1038/s41586-019-1923-7 (2020).

9 Dauparas, J. et al. Robust deep learning-based protein sequence design using ProteinMPNN. Science 378, 49–56, doi:10.1126/science.add2187 (2022).

10 Hsu, C. et al. Learning inverse folding from millions of predicted structures. 2022.2004.2010.487779, doi:10.1101/2022.04.10.487779 %J bioRxiv (2022).

11 Akpinaroglu, D. et al. Structure-conditioned masked language models for protein sequence design generalize beyond the native sequence space. 2023.2012.2015.571823, doi:10.1101/2023.12.15.571823 %J bioRxiv (2023).

12 Gao, Z., Tan, C., Chacón, P. & Li, S. Z. J. a. p. a. PiFold: Toward effective and efficient protein inverse folding. (2022).

13 O’Connell, J. et al. SPIN2: Predicting sequence profiles from protein structures using deep neural networks. Proteins 86, 629–633, doi:10.1002/prot.25489 (2018).

14 Ingraham, J., Garg, V. K., Barzilay, R. & Jaakkola, T. Generative models for graph-based protein design. Advances in Neural Information Processing Systems 32 (Nips 2019) 32 (2019).

15 Strokach, A., Becerra, D., Corbi-Verge, C., Perez-Riba, A. & Kim, P. M. Fast and Flexible Protein Design Using Deep Graph Neural Networks. Cell Systems 11, 402-+, doi:10.1016/j.cels.2020.08.016 (2020).

16 Mu, J. X. et al. Graphormer supervised de novo protein design method and function validation. Briefings in Bioinformatics 25, doi:10.1093/bib/bbae135 (2024).

17 Zhang, Y. et al. ProDCoNN: Protein design using a convolutional neural network. Proteins-Structure Function and Bioinformatics 88, 819–829, doi:10.1002/prot.25868 (2020).

18 Notin, P. et al. ProteinGym: Large-Scale Benchmarks for Protein Design and Fitness Prediction. bioRxiv, doi:10.1101/2023.12.07.570727 (2023).

19 Gao, Z. et al. Proteininvbench: Benchmarking protein inverse folding on diverse tasks, models, and metrics. 36 (2024).

20 Johnson, S. R. et al. Computational scoring and experimental evaluation of enzymes generated by neural networks. 1–10 (2024).

21 Yu, J., Mu, J., Wei, T. & Chen, H. F. Multi-indicator comparative evaluation for deep learning-based protein sequence design methods. Bioinformatics 40, doi:10.1093/bioinformatics/btae037 (2024).

22 Cao, Y., Geddes, T. A., Yang, J. Y. H. & Yang, P. Y. Ensemble deep learning in bioinformatics. Nat Mach Intell 2, 500–508, doi:10.1038/s42256-020-0217-y (2020).

23 Xu, S. & Onoda, A. Accurate and Fast Prediction of Intrinsically Disordered Protein by Multiple Protein Language Models and Ensemble Learning. J Chem Inf Model 64, 2901–2911, doi:10.1021/acs.jcim.3c01202 (2024).

24 Wee, J. & Xia, K. Persistent spectral based ensemble learning (PerSpect-EL) for protein-protein binding affinity prediction. Brief Bioinform 23, doi:10.1093/bib/bbac024 (2022).

25 Qu, Y. et al. Ensemble Learning with Supervised Methods Based on Large-Scale Protein Language Models for Protein Mutation Effects Prediction. Int J Mol Sci 24, doi:10.3390/ijms242216496 (2023).

26 Gao, Z., Tan, C. & Li, S. Z. J. a. p. a. Alphadesign: A graph protein design method and benchmark on alphafolddb. (2022).

27 Zheng, Z. et al. in International conference on machine learning. 42317–42338 (PMLR).

28 Henikoff, S. & Henikoff, J. G. Amino acid substitution matrices from protein blocks. Proc Natl Acad Sci U S A 89, 10915–10919, doi:10.1073/pnas.89.22.10915 (1992).

29 Llinares-Lopez, F., Berthet, Q., Blondel, M., Teboul, O. & Vert, J. P. Deep embedding and alignment of protein sequences. Nat Methods 20, 104–111, doi:10.1038/s41592-022-01700-2 (2023).

30 Lin, Z. M. et al. Evolutionary-scale prediction of atomic-level protein structure with a language model. Science 379, 1123–1130, doi:10.1126/science.ade2574 (2023).

31 Yang, P. et al. Allele-Specific Suppression of Variant MHC With High-Precision RNA Nuclease CRISPR-Cas13d Prevents Hypertrophic Cardiomyopathy. Circulation 150, 283–298, doi:10.1161/CIRCULATIONAHA.123.067890 (2024).

32 Minarik, P., Tomaskova, N., Kollarova, M. & Antalik, M. Malate dehydrogenases--structure and function. Gen Physiol Biophys 21, 257–265 (2002).

33 Repecka, D. et al. Expanding functional protein sequence spaces using generative adversarial networks. 3, 324–333 (2021).

34 Qiu, J. et al. InstructPLM: Aligning Protein Language Models to Follow Protein Structure Instructions. 2024.2004.2017.589642, doi:10.1101/2024.04.17.589642 %J bioRxiv (2024).

35 Chu, A. E., Lu, T. Y. & Huang, P. S. Sparks of function by de novo protein design. Nat Biotechnol 42, 203–215, doi:10.1038/s41587-024-02133-2 (2024).

36 Listov, D., Goverde, C. A., Correia, B. E. & Fleishman, S. J. Opportunities and challenges in design and optimization of protein function. Nat Rev Mol Cell Bio 25, 639–653, doi:10.1038/s41580-024-00718-y (2024).

37 Khakzad, H. et al. A new age in protein design empowered by deep learning. Cell Syst 14, 925–939, doi:10.1016/j.cels.2023.10.006 (2023).

38 Pillai, A. et al. De novo design of allosterically switchable protein assemblies. Nature 632, 911–920, doi:10.1038/s41586-024-07813-2 (2024).

39 Ganaie, M. A., Hu, M., Malik, A. K., Tanveer, M. & Suganthan, P. N. J. E. A. o. A. I. Ensemble deep learning: A review. 115, 105151 (2022).

40 Sagi, O., Rokach, L.J.W.i.r.d.m. & discovery, k. Ensemble learning: A survey. 8, e1249 (2018).

41 Madani, A. et al. Large language models generate functional protein sequences across diverse families. Nat Biotechnol 41, 1099–1106, doi:10.1038/s41587-022-01618-2 (2023).

42 Nijkamp, E., Ruffolo, J. A., Weinstein, E. N., Naik, N. & Madani, A. ProGen2: Exploring the boundaries of protein language models. Cell Syst 14, 968–978 e963, doi:10.1016/j.cels.2023.10.002 (2023).

43 Watson, J. L. et al. De novo design of protein structure and function with RFdiffusion. Nature 620, 1089–1100, doi:10.1038/s41586-023-06415-8 (2023).

44 Dai, D. et al. Deepseekmoe: Towards ultimate expert specialization in mixture-of-experts language models. (2024).

45 Cai, W. et al. A survey on mixture of experts. (2024).

46 Vaswani, A. et al. Attention Is All You Need. Adv Neur In 30 (2017).

47 Kabsch, W. & Sander, C. Dictionary of protein secondary structure: pattern recognition of hydrogen-bonded and geometrical features. Biopolymers 22, 2577–2637, doi:10.1002/bip.360221211 (1983).

48 Ben Chorin, A. et al. ConSurf-DB: An accessible repository for the evolutionary conservation patterns of the majority of PDB proteins. Protein Sci 29, 258–267, doi:10.1002/pro.3779 (2020).

